# Peptides derived from plant virus VPg protein inhibit eIF4E oncogene

**DOI:** 10.1101/298455

**Authors:** Izabela Wojtal, Malgorzata Podsiadla-Bialoskorska, Renata Grzela, Malgorzata Bujak, Ewa Szolajska, Jadwiga Chroboczek

## Abstract

Viruses of the *Potyviridae* family have VPg protein covalently attached to the 5’ end of their linear RNA genome. The protein interacts with the host translation initiation factor eIF4E that occurs in plant cells in two isoforms, one being the preferable target of a given VPg, the remaining one still acting in host protein synthesis. In animal cells only one form of eIF4E is directly involved in protein synthesis. The human eIF4E is known to be an oncogene; elevated expression of eIF4E leads to oncogenic transformation, cancers in animal models and poor prognosis in human cancers, while reduction of the eIF4E level can reverse the transformed phenotype. We show that VPg protein delivery to cells containing only one eIF4E isoform involved in protein synthesis resulted in immobilization of eIF4E in the cytoplasm. The region of VPg involved in the interaction with eIF4E has been partially identified. Peptides derived from this region interacted better with eIF4E than complete VPg protein. Here we characterized one of VPg peptides, VPg5 and we show that VPg5 delivered to colon carcinoma HCT116 cells is able to inhibit cell growth, which is accompanied by reduction in eIF4E level.

**List of abbreviations:**

AcMNPV: Autographa californica multicapsid nucleopolyhedrovirus
CBB: Coomassie Brilliant Blue
ClYVV: Clover yellow vein virus
eIF4E: eukaryotic initiation translation factor 4E
4E: BP, eIF4E binding protein
FCS: fetal calf serum
LMV: Lettuce Mosaic Virus
MOI: multiplicity of infection
MW: molecular weight
ON: overnight
TMB: 3,3,5,5-tetramethylbenzidine
*T. ni*: *Trichoplusia ni* cells
PBST: PBS buffer containing 0.05% Tween-20
pi: post infection
PI: propidium iodide
TEV: tobacco etch virus
TVMV: tobacco vein mottling virus
PVY: potato virus Y
VPg: genome-linked viral protein

## Introduction

Potyvirus VPg is a multifunctional protein implicated in translation, long-distance movement, replication and suppression of host antiviral silencing response (1, 2). The host partner of VPg is the eIF4E, an mRNA 5‘ cap-binding protein. The eIF4E and VPg are the major determinants of resistance *versus* susceptibility to potyvirus infection; plants are virus-resistant when mutations in eIF4E or in VPg preclude their interaction (3–6). However, higher plants produce two isoforms of eIF4E and potyviral VPgs display greater affinity to one or the other eIF4E isoform (7, 8). Thus, although host protein synthesis is weakened upon infection, the virus cell cycle can be completed since the second eIF4E remains active in plant metabolism necessary for virus development.

In human cells only one cap-binding protein, eIF4E-1, directly promotes protein synthesis (9), and a significant fraction of it (sometimes up to 70%) is found in the nucleus (10). In the cytoplasm, eIF4E is involved in the initiation of protein synthesis while in the nucleus it is implicated in transport of certain mRNAs to the cytoplasm. The nuclear eIF4E has an affinity for a class of mRNAs that specify the short-living proteins implicated in cell cycle progression and malignant transformation (11). An unchecked overexpression of eIF4E can cause cell growth and malignant transformation in tissue cultures and is a feature of experimental cancers, whereas reduction of the eIF4E level can reverse the transformed phenotype (12, 13). Elevated expression of eIF4E in human cancers correlates with poor prognosis (11, 14). These effects can partly be explained by a higher level of short-lived proteins involved in malignant transformation, stemming from the increased eIF4E-dependent transport of the appropriate mRNAs to the cytoplasm. The proof of hypothesis was furnished by the human trial in which the inhibition of eIF4E activity in acute myeloid leukemia patients brought about quite positive results, whereby cancer remission was associated with eIF4E delocalization from the nucleus to the cytoplasm (15). It appears that eIF4E is an oncogene that acts as a node governing proliferation and survival signaling (16–18).

We have shown that Potato Virus Y (PVY) VPg interacts *in vitro* with various eIF4Es, including animal eIF4E (19). It was thus interesting to study what will happen in the presence of VPg in cells with only one form of the eIF4E factor active in protein synthesis. We observed that in the presence of VPg the nuclear pool of eIF4E was dramatically depleted since VPg immobilized initiation factor in the cytoplasm. Recently, we partially characterized the site of PVY VPg responsible for interaction with eIF4E, both of plant and human origin, and constructed peptides derived from this region, some of which interacted better with eIF4E than the complete VPg protein. We wished to show that one of VPg-derived peptides is able inhibit the pro-oncogenic role of eIF4E in neoplastic cells. We have chosen peptide VPg5 for further work and also prepared VPg5 mutants interacting weakly with human eIF4E. Delivery of VPg5 to neoplastic cells in culture resulted in cell death accompanied by eIF4E reduction.

## Results

### Localization of eIF4E and VPg in insect cells

Upon infection with the recombinant baculovirus expressing PVY VPg, the VPg protein was observed only in the cytoplasm (Fig. 1A, VPg column). The endogenous initiation factor eIF4E localized at the beginning of infection to both, the nucleus and the cytoplasm, but at 60 h pi. was observed only at the cell periphery (Fig. 1A, eIF4E column). Indeed, at the beginning of infection, a significant fraction of the eIF4E was in the nucleus but diminished in the course of infection; at 60 h pi. eIF4E was observed in the cytoplasm only, showing significant overlap with VPg. To find out if the observed changes in the eIF4E localization were due to the presence of VPg, we analyzed the localization of eIF4E upon infection with empty bacmid and with baculovirus expressing either a non-relevant protein (the domain II of HCV helicase) or VPg (Fig. 1B). At 60 h pi, only cells expressing VPg displayed eIF4E disappearance from the nucleus (Fig. 1B, lowest panel), demonstrating the specificity of VPg effect. To obtain more quantitative data, after infection with the recombinant baculovirus expressing VPg the insect cell nuclei were separated from the cytoplasm, and eIF4E level in both compartments was assessed by Western blot (Fig. 1C) followed by densitometry (Table 1). At 12h a comparable level of eIF4E was present in the cytoplasm and in the nucleus. The nuclear pool of the eIF4E diminished somewhat in the course of infection with bacmid and with the helicase-expressing virus, from 51-53% down to 37-40% (Table 1). However, upon expression of his-tagged or non-tagged VPg forms, the nuclear pool of the eIF4E was much more severely depleted; at 24 and 36h pi it accounted for only 13-18% of total cellular factor. Interestingly, VPg expression was significantly lower than that of HCV helicase, which underlies the strength of VPg effect (Fig. 1C, lower panel on the right).

**Table 1.**
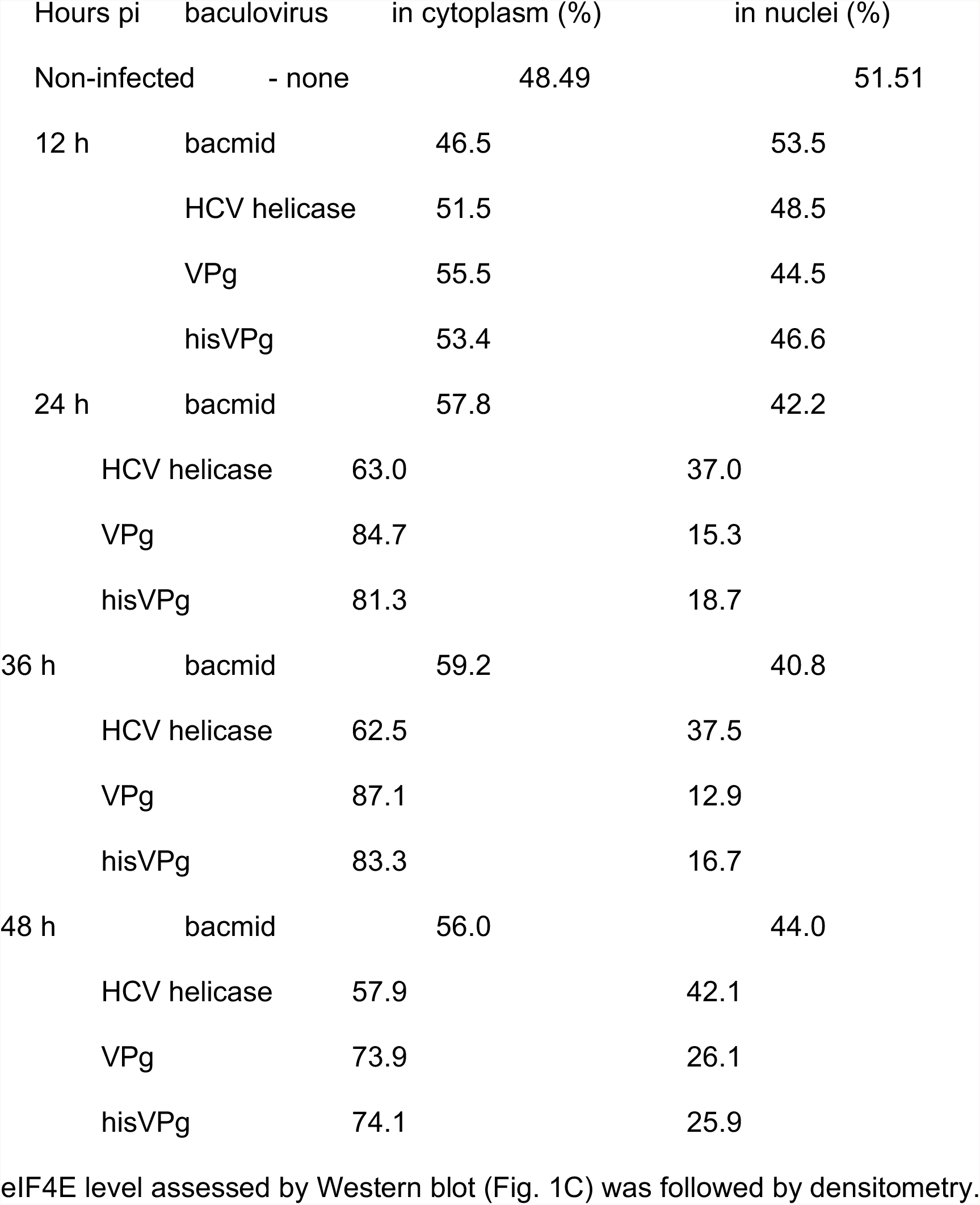
Distribution of eIF4E in insect cells

| Hours pi | baculovirus | in cytoplasm (%) | in nuclei (%) |
| --- | --- | --- | --- |
| Non-infected | - none | 48.49 | 51.51 |
| 12 h | bacmid | 46.5 | 53.5 |
|  | HCV helicase | 51.5 | 48.5 |
|  | VPg | 55.5 | 44.5 |
|  | hisVPg | 53.4 | 46.6 |
| 24 h | bacmid | 57.8 | 42.2 |
|  | HCV helicase | 63.0 | 37.0 |
|  | VPg | 84.7 | 15.3 |
|  | hisVPg | 81.3 | 18.7 |
| 36 h | bacmid | 59.2 | 40.8 |
|  | HCV helicase | 62.5 | 37.5 |
|  | VPg | 87.1 | 12.9 |
|  | hisVPg | 83.3 | 16.7 |
| 48 h | bacmid | 56.0 | 44.0 |
|  | HCV helicase | 57.9 | 42.1 |
|  | VPg | 73.9 | 26.1 |
|  | hisVPg | 74.1 | 25.9 |

eIF4E level assessed by Western blot (Fig. 1C) was followed by densitometry.

**Fig. 1.**
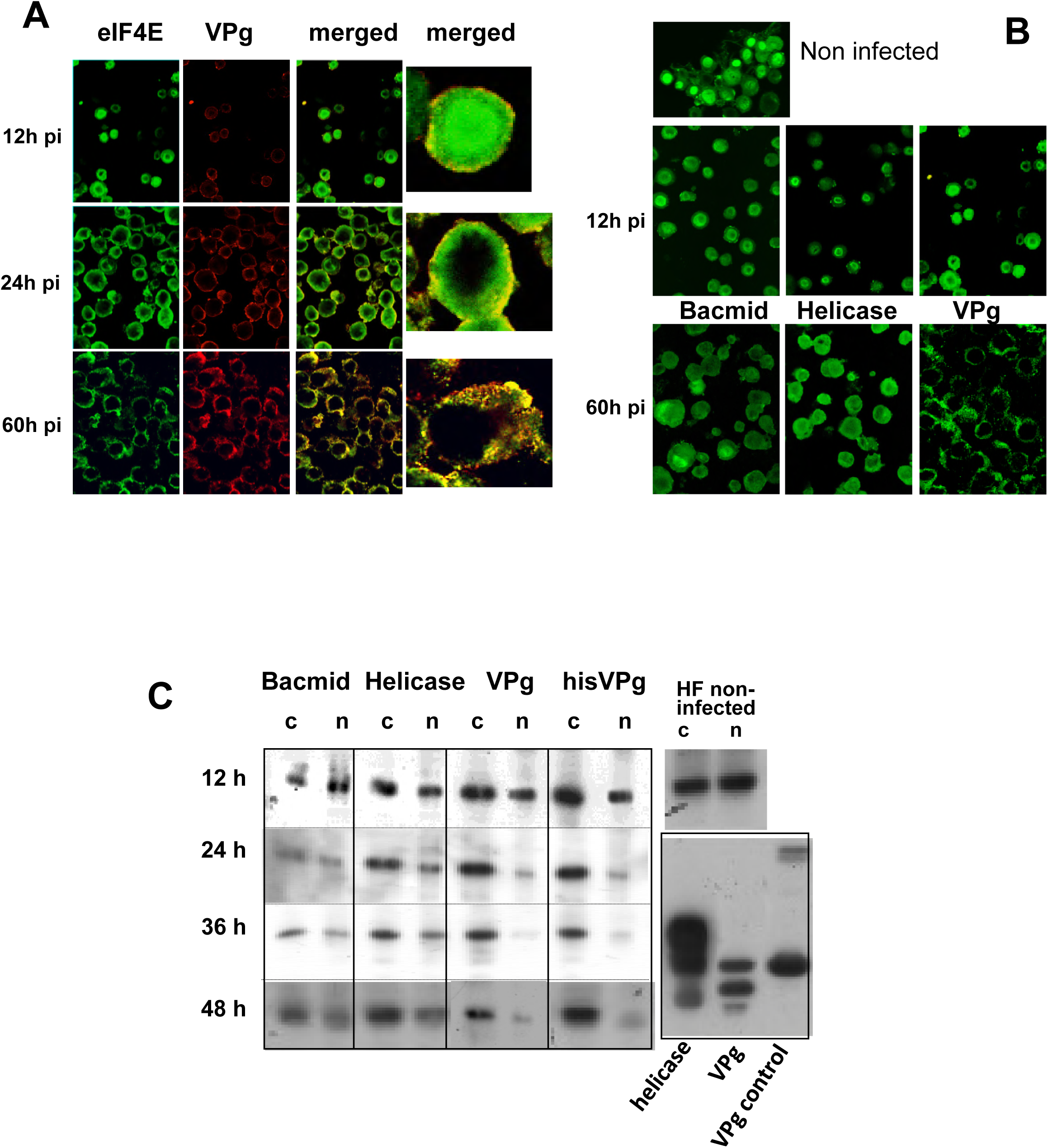
Effect of VPg on distribution of endogenous eIF4E in insect cells. Expressing insect cells were analyzed by confocal microscopy (× 60). (A) Localization of eIF4E (in green) in the presence of VPg (in red). (B) Cytoplasmic retention of native eIF4E (in green) is specific for VPg. (C) Distribution of endogenous insect eIF4E upon VPg expression (MOI 10). Expressing cells nuclei (n) were separated from the cytoplasm (c) and the eIF4E content was analyzed. Recombinant proteins expression is shown in the lower far right panel.

These experiments demonstrated that (1) about half of the total cellular eIF4E synthesized in the cytoplasm is transported into the nucleus, confirming results seen for other eukaryotic cells (10); (2) VPg expressed alone in insect cells, despite carrying the putative nuclear localization signal, is localized in the cytoplasm only; (3) in the presence of VPg, eIF4E is retained in the cytoplasm, which results in depletion of the nuclear pool of eIF4E.

### Interaction of VPg peptides with eIF4E

Previously, upon selection of proteolytically cleaved VPg protein on a column-immobilized wheat eIF(*iso*)4E, we tentatively identified an eIF4E-interacting site on VPg (19). These results suggested that the N-terminal part of the VPg up to tyrosine 40 is dispensable for the interaction, and that interaction site involves downstream region, starting with arginine 41. We have constructed five peptides derived from the putative interaction site and expressed them in frame with the N-terminally attached GST protein, expressing similarly also a complete VPg protein. These proteins after purification have been used in the ELISA sandwich assay, in which peptide attached to eIF4E was recognized with the anti-GST antibody (Fig. 2). The interaction with wheat eIF(*iso*)4E and with human eIF4E were analysed. The best interacting peptides were VPg5 and VPg4 for the plant factor. The human factor interacted best with the same peptides and in addition with the peptide VPg1. This shortest interacting peptide (VPg1, 19 aa) was contained in both VPg5 (26 aa) and VPg4 (42 aa). Also, the VPg2 contained peptide VPg1 but was significantly longer and much more hydrophobic in character. It attached less well to eIF4E and, similarly as peptide VPg3, had a propensity for forming oligomers at high concentration (results not shown). We have chosen to work with peptide VPg5. In parallel we created two VPg5 mutants. Based on comparison of corresponding sequences from other potyviral VPgs (Fig. 3A), we changed either tyrosine 64 to alanine (pVPg5A1) or tyrosine 64 and glycine 54 to alanines (pVPg5A2). These peptides in fusion with GST were expressed and purified (Fig. 3C). The VPg5 mutants were analysed for their interaction with human eIF4E (Fig. 3D). Both VPg5 mutants interact rather weakly with human eIF4E, only slightly better than with GST. Interestingly, rather weak interaction could be observed also for a complete VPg protein (but which was on the same level as that shown in Fig. 2).

**Fig. 2.**
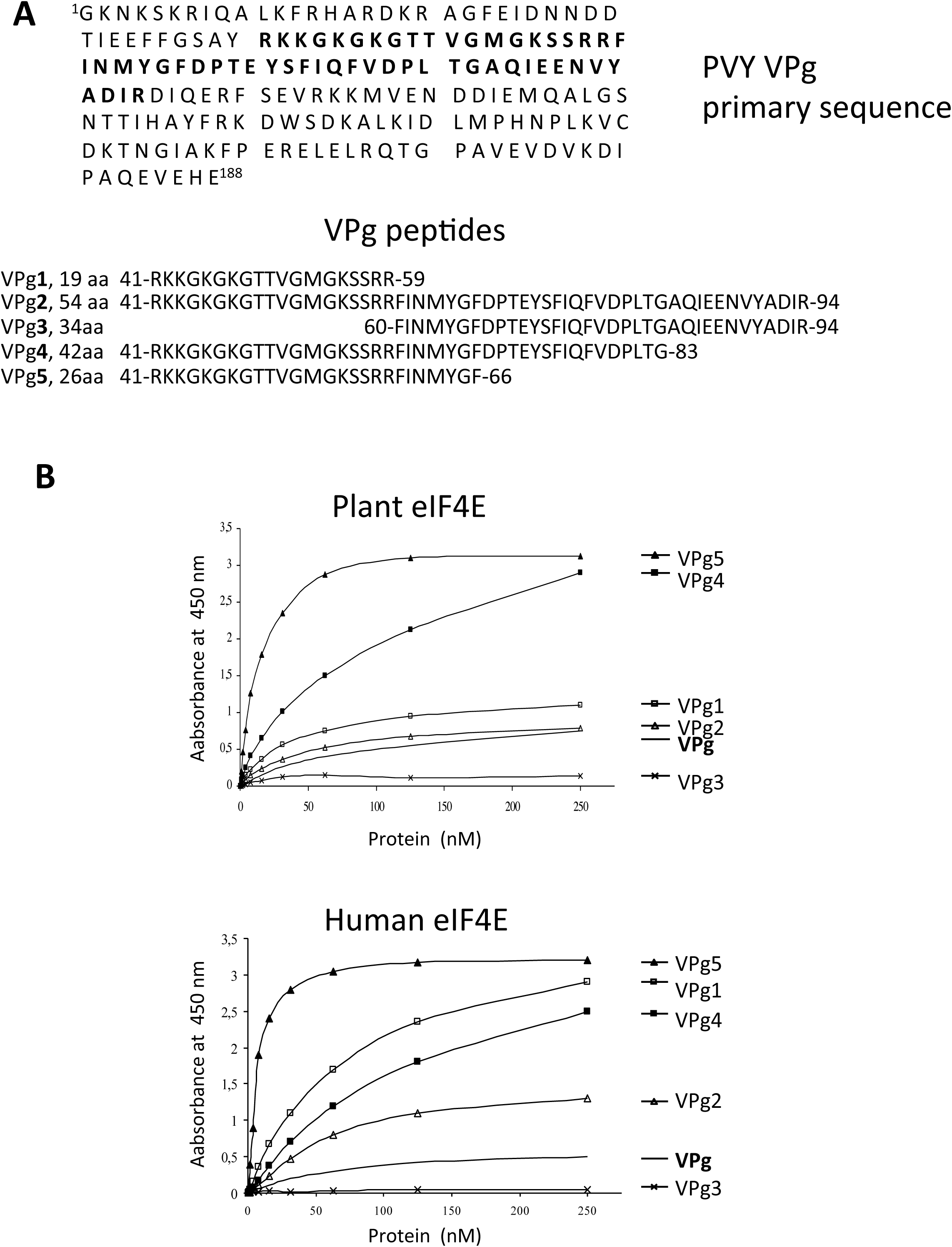
Interaction of VPg peptides with eIF4E. (A) Amino acid sequence of VPg and VPg peptides derived from the VPg region involved in interaction with eIF4E. The putative eIF4E interaction site is marked with bold letter. (B) Interaction of VPg peptides with wheat eIF(*iso*)4E and human eIF4E. Complete VPg protein is marked in bold.

**Fig. 3.**
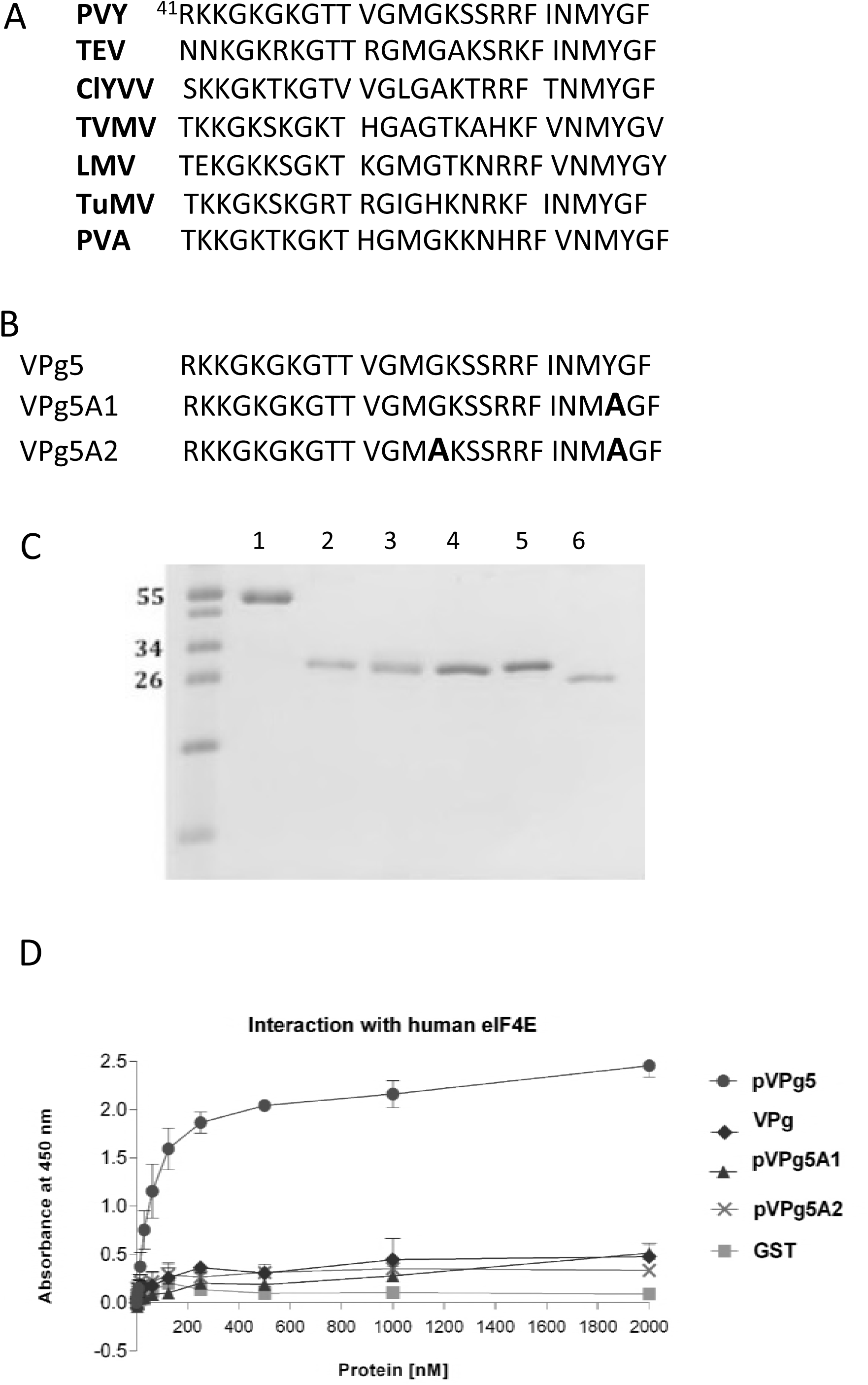
VPg5 peptides and their interaction with human eIF4E. (A) Comparison of primary sequence of some potyviral VPgs. (B) Primary sequence of VPg5 peptides. (C) GST-fusion proteins used in ELISA test, run on 15% tricine polyacrylamide gel (41). Samples of 1-2 µg were stained with CBB. Lane 1 – VPg protein, lane 2 – VPg5; lane 3 - VPg5A1; lane 4 - VPg5A2; lane 5 – his-eIF4E protein; lane 6 - GST protein. (D) Interaction of pVPg peptides with human eIF4E (ELISA sandwich technique).

### Effect of VPg peptides on neoplastic cells in culture

This effect was investigated on HCT116 colon carcinoma cells in culture, transduced with Tat fusions of VPg5 and VPg5A2. Treatment of neoplastic cells with TAT-VPg5 resulted in marked cell death, while TAT-VPgA2 despite low binding to eIF4E appears to act only somewhat less well than Tat-VPg5 (Fig. 4). Cell death induced by Tat-VPg5 was accompanied by reduction in eIF4E level (Fig.4, BC).

**Fig. 4.**
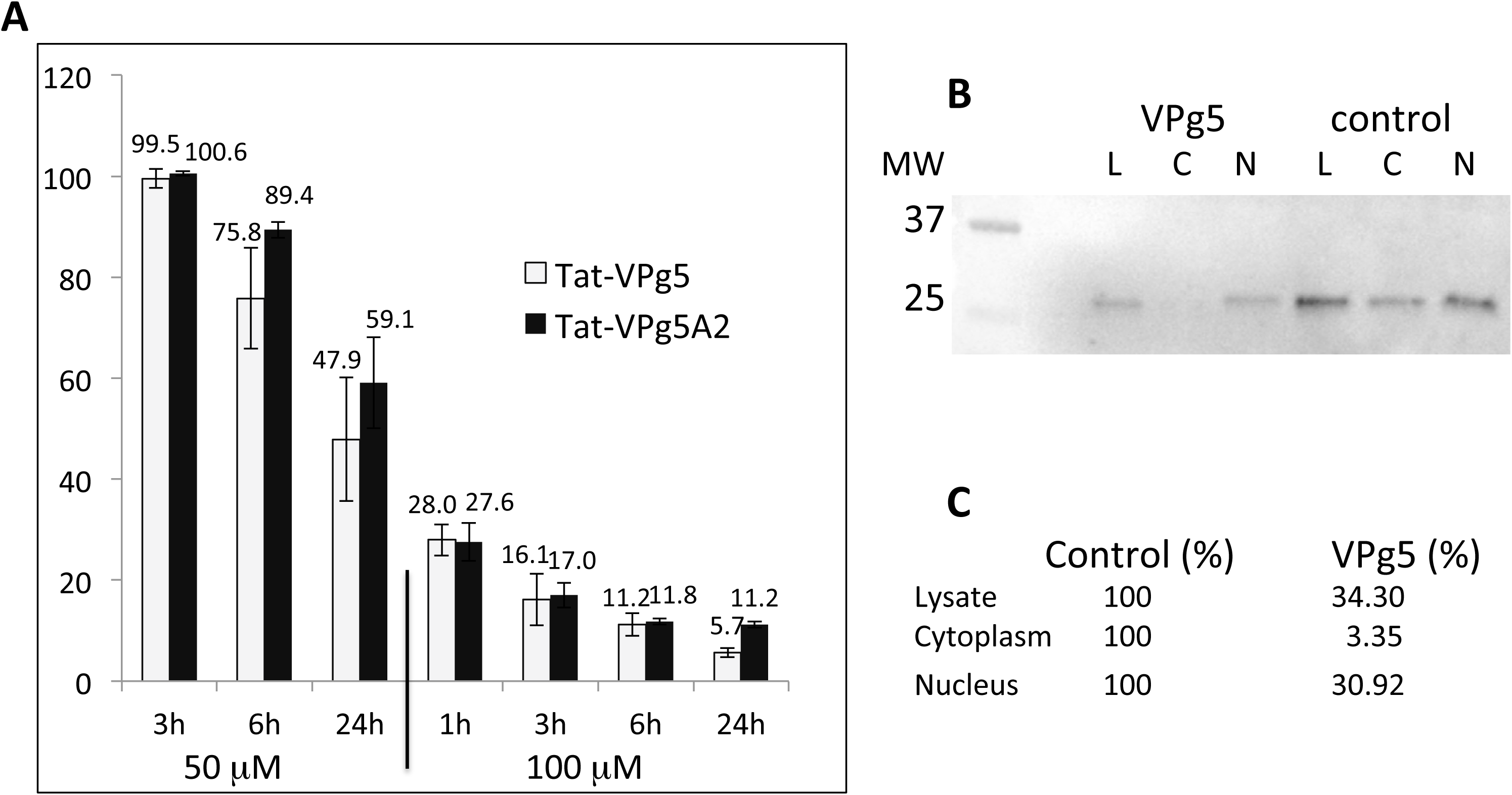
Effect of Tat-VPg peptides on HCT116 cell viability. Tat-VPg5 peptides were applied to HCT116 cells as described in Materials and Methods. (A). Cell viability was measured by trypan blue exclusion. B. Portions of 100 000 cells untreated or treated with Tat-VPg5 underwent fractionated with REAP method (40), run on SDS-PAGE gel, and Western blot was performed with anti-eIF4E antibody. Densitometry was performed as described in Materials and Methodes.

## Discussion

Nuclear eIF4E participates in a variety of important RNA processing events including nucleocytoplasmic transport of specific, growth regulatory mRNAs. Indeed, the availability of eIF4E is believed to be critical for nuclear export and translational activation of capped mRNAs with extensive secondary structures in their 5’ untranslated regions (UTRs) (20), many of which code for the labile regulatory proteins essential for cell growth or viability. Various molecular mechanisms of cell growth converge on eIF4E, and it appears that eIF4E is a critical node in RNA regulation influencing nearly every stage of cell cycle progression (16). Thus, eIF4E, besides its direct cytoplasmic role in protein synthesis, seems to have another, nuclear function, whereby by promoting the export of messenger RNA of several proteins involved in cell cycle it plays a key role in regulation of cell growth and survival.

Elevated expression of eIF4E leads to oncogenic transformation in tissue culture, cancers in animal models and poor prognosis in some human cancers. Reduction of eIF4E level can reverse the transformed phenotype, which suggests that eIF4E could be exploited as a therapeutic target. A validation of eIF4E as a therapeutic target has been provided by *in vivo* experiments, in which suppression of eIF4E expression in mice by eIF4E-specific antisense oligonucleotides (ASO) reduced tumor growth in mice (21). Despite reducing the eIF4E levels in mouse liver by 80%, administration of eIF4E-specific ASO did not affect the body weight or the liver enzymes. This suggested that the malignant tumors might be more susceptible to eIF4E inhibition than the normal tissues. Most importantly, a human trial on acute myeloid leukemia patients in whom eIF4E was inactivated with ribavirin, a cap structure mimic (and thus an eIF4E inhibitor) resulted in tumor growth inhibition and cancer remission that were correlated with eIF4E reduction and translocation of nuclear eIF4E to the cytoplasm in blasts (15, 22). Recently, a phase I trial of ribavirin and low-dose cytarabine for the treatment of acute myeloid leukemia with elevated eIF4E has been terminated, showing that the combination was well tolerated, with some patients having a full molecular response with both reduction of eIF4E levels and eIF4E re-localization to cytoplasm (23).

The eIF4E binding partners such as 4E-BPs, Gemin5, Xenopus Maskin and mammalian neuroguidin, have all been shown to exert translational control (24). These factors usually block the interaction between eIF4E and the scaffold-like eIF4G factor, involved in the formation of the large translation initiation complex eIF4F (25). They all contain the dorsal eIF4E-binding motif YxxxxLØ. Another type of eIF4E effectors, the PML protein and the arenaviral protein Z use their respective RING domains to directly interact with the dorsal surface of eIF4E, thus reducing the affinity of eIF4E for its ligand, 5’ cap of mRNA (26, 27). These two proteins shuttle between nucleus and cytoplasm (28), and thus can interact with both nuclear and cytoplasmic eIF4E pools. In addition, a transcription factor, homeodomain protein PRH/HEX that binds nuclear eIF4E inhibits eIF4E function only when its NLS is intact, i.e. it must be nuclear to inhibit eIF4E (29), whereas VPg protein delivered alone shows an exclusively cytoplasmic localization. More importantly, VPg does not contain the eIF4E-binding motif YxxxxLØ or the RING domain, which suggests that VPg interaction with eIF4E occurs at different eIF4E site. Some plants became resistant to potyviruses due to the eIF4E mutations abolishing interaction with VPg. Such mutations have been identified in two eIF4E regions; one partially overlapping the interior of the cap-binding pocket (which is in the concave part of eIF4E) and another on the surface of eIF4E facing 90° from the cap-binding site (30, 7). Both these regions are distant from the dorsal eIF4E site attaching YxxxxLØ peptides.

Another member of eIF4E family of translational initiation factors that interact with the 5‘cap structure of mRNA is 4E-HP. However, 4E-HP does not share the role of eIF4E in mRNA transport from nucleus to cytoplasm; it is found exclusively in the cytoplasm whereby it may act as a translational repressor of mRNAs (31). The third member of the eIF4E family, eIF4E3, that share ∼25% identity and ∼50% similarity with the two other eIF4E family members, and is nuclear and cytoplasmic, also binds the m7G cap, but acts as a tumor suppressor, competing with the growth-promoting functions of eIF4E (18). All these observations together show that VPg protein is novel type of eIF4E inhibitor.

Different type of eIF4E inhibitor in a form of a cap analog was used by us during *in vivo* studies in the animal model of hepatocellular carcinoma. HCC is one of the leading causes of cancer-related death worldwide, one of the most lethal, chemoresistant and prevalent cancers in the human population with limited treatment options (32). It has been shown that HCC development depended on eIF4E overexpression (33). We have demonstrated that eIF4E inhibition with cap analog in the orthotopic rat model of HCC resulted in tumor growth inhibition, accompanied by decrease in the level of two pro-oncogenes, eIF4E and c-myc (34). Of nWojtl et al, Mrch 2018ote, c-myc is amplified in up to 50% of HCC cases (35).

This time we used a peptide-type inhibitor of eIF4E to find out whether inhibition of the eIF4E activity is an interesting therapeutic approach to cancer treatment that conceivably could be extended to other neoplasms driven by eIF4E overexpression. A partially identified site on PVY VPg served here as a source of five peptides, out of which some interacted better with eIF4E than complete VPg protein. We have chosen a peptide VPg5 of 26 amino acid residues that strongly interacted with eIF4E (Figs. 2 and 3). In parallel we designed a mutant VPg5A2, which showed rather low-level interaction with eIF4E. These peptides were delivered to neoplastic HCT116 cells in a form of the N-terminal fusions with transducing Tat peptide (36). As use of Tat peptide might introduce several artefacts, we followed the conditions developed for minimizing the nonspecific Tat peptide activity (37). Viability of neoplastic cells was significantly diminished by addition of Tat-VPg5 peptides, in a dose dependent manner, quite strongly with the original VPg5 peptide, somewhat less with VPg5A2 peptide. Of note, designing the control - eIF4E-nonreacting - peptide is rather difficult, as there is no structural data identifying the amino acid residues of PVY VPg interacting with eIF4E.

It is possible that NLS of VPg (probably starting from ^41^Arg) is not functioning when the complete protein (188 amino acid residues) is delivered upon expression in insect cells, as complete VPg brought alone was localized exclusively to cytoplasm (Fig. 1). However, it is quite plausible to think that NLS in the VPg5 peptide could be functional and VPg5 would go directly to the nucleus, interacting there with the pro-oncogenic nuclear eIF4E. Nevertheless, the eIF4E localization seen after treatment with VPg5 (Fig. 4 B,C) shows strong reduction of nuclear eIF4E but nearly complete depletion of factor in the cytoplasm. Further experiments including confocal microscopy would more precisely describe the intracellular effect and localization of VPg5.

In conclusion, results obtained with VPg5 peptide show its activity as a novel, efficient inhibitor of eIF4E, leading to death of neoplastic cells, which is accompanied by reduction in eIF4E level. This kind of specific oncogene eIF4E inhibitors should be analysed in eIF4E-driven cancers; it was thought that about 30% of human malignancies show overexpression of eIF4E (38), however, more detailed data indicate this percent as close to 63% (39).

## Materials and methods

### Materials

*T. ni* insect cells (Invitrogen) were grown as described by Grzela et al (19). Anti-eIF4E mAb that recognized human and animal eIF4E was from Santa Cruz Biotechnology (USA). Anti-eIF4E recognizing human, mouse and insect eIF4E, was a gift of Nahum Sonenberg (McGill University, Canada). Anti-VPg rabbit polyclonal antibody was prepared in ESD (France), using as antigen PVY hisVPg expressed in baculovirus (19) and was affinity-purified with VPg immobilized on CNBr-activated Sepharose 4B (Amersham Bioscience). Goat anti-GST-HRP antibody was from Rockland Immunochemicals Inc. Molecular weight markers for SDS-PAGE were from Thermo Scientific. Rabbit polyvalent anti-Dd serum was prepared in the laboratory using as antigen a mixture of well-formed and ultrasound-treated Dd.

### Localization of eIF4E and VPg in insect cells

*T. ni* cells (2×10^6^ cells/ml) were infected at MOI 10 with recombinant baculoviruses or empty bacmid. Cells were collected at indicated times p.i., washed with PBS, fixed in cold 2% PFA and suspended in 300 µl PBS. Portions of 100 µl of fixed cells on round coverslips inserted into wells of a 24-wells plate and rehydrated were permeabilized with cold 0.1% Trition X-100/PBS, rinsed twice with PBS and blocked with 5% serum/PBS for 30 min at RT. Permeabilized cells were incubated for 1 h with anti-his or anti-eIF4E Ab diluted 100-fold, washed and incubated for 1 h with anti-rabbit FITC-conjugated Ab or anti-mouse Texas Red-conjugated Ab (Jackson ImmunoResearch), diluted 1:100. Nuclei were stained with PI in PBS (1 µg/ml) and PBS-rinsed coverslips were mounted with 50% glycerol. Images were collected with Bio-Rad MRC-600 laser scanning confocal apparatus coupled to a Nikon Optiphot microscope.

### Insect cell fractionation

High Five cells (2×10^6^ cells/ml) were infected with the baculoviruses expressing VPg, or hisVPg, or domain II of HCV helicase or empty bacmid. The infected cells were collected at the indicated times p.i. by centrifugations at 2000 rpm for 10 min and nuclei were separated from the soluble fraction as described (Xu et al, 1995). Cytoplasmic and nuclear fractions were run on 15% SDS-PAGE followed by Western blot with anti-eIF4E antibody and the ECL system. Densitometry of eIF4E bands was performed with GelDoc 2000 (BioRad, Quantity One software).

### Protein expression and purification

VPg expression in *E. coli* and purification was done according Grzela et al (19). GST protein was expressed in *E. coli* BL21(DE3) Rosetta from pGEX-4TI and purified on the GST-Sepharose. For the synthesis of the fusion protein GST-VPg5, the VPg5 gene encoding a part of VPg protein between residues 41 – 66, was amplified by PCR using plasmid pGEXVPg (containing the gene encoding the VPg protein of PVY strain 0, acc. no Z29526; Grzela *et al*., 2006) as template, with the forward primer 5’CGCGGATCCGAAAACCTGTATTTTCAGGGCAGGAAAAAGGGAAAAGGTAAA GG3’ and the reverse primer 5’CCGCCGCTCGAGCTATTAGAACCCGTACATGTTAATGAA3’. The resulting PCR product was then digested by BamHI/XhoI and inserted into pGEX4T-1 vector between BamHI and XhoI sites for expression in *E. coli* as GST-fusion protein.

Two VPg5 mutants, VPg5A1 and VPg5A2 have been prepared in form of a fusion with the GST protein (at the N-terminus of the peptide). To obtain GST-VPg5A2, mutagenesis was done using Q5^®^ Site-Directed Mutagenesis Kit (NEB) and a GST-VPg5 in pGEX4T-1 as a template. In VPg5A1 the Gly54 has been changed to Ala, in VPg5A2 both Gly54 and Tyr64 has been changed to Ala. The following primers (forward and reverse, F and R) have been used for mutagenesis of the pVPg5 sequence. The correct sequence of mutants was confirmed by DNA sequencing.

VPg5A1

F1A AGCAGGAGGTTCATTAACATGGCCGGGTTCTAATAGCTCGAG

R1A TGACTTGCCCATACCAACTGTGGTACCTTTACC

VPg5A2

F2A

AAAGGTACCACAGTTGGTATGGCCAAGTCAAGCAGGAGGTTCATTAACATGGCCGGGTTCTAATAG

R2A

ACCTTTTCCCTTTTTCCTGCCCTGAAAATACAGGTTTTCGGATCCACGCGGAACCAG

GST peptides were expressed in *E. coli* Rosetta2(DE3)pLyS as follows. Bacteria were grown in the presence of 100 μg/ml of carbenicilillin and 20 μg/ml chloramphenicol, initially at 37°C. Protein production was induced with 400 μM IPTG at OD_600_ equal 0.6, then the temperature was switched to 30^°^C and 4h later the bacteria were harvested. Cell pellet from 250 ml culture was suspended in 10 ml of 50 mM Hepes buffer, pH 7.4, containing 300 mM NaCl, 5 mM DTT, 0.5 mM EDTA and supplemented with the protease inhibitor cocktail (Complete, Roche) and lysozyme (LabEmpire) at 1mg/ml. The bacteria were lysed by sonication on ice. The lysates were then centrifuged at 14 000 rpm for 30 min at 4°C and the supernatants were filtered through 0.2 μm filter before applying onto 0.5 ml Glutathione-Sepharose 4B column (GE Healthcare). After 4 washes with 10 ml of lysis buffer GST-fusion proteins were eluted with five 1ml portions of 10 mM reduced glutathione in lysis buffer.

Human eIF4E was cloned in plasmid pETM30 (EMBL) with 6xhis tag attached at the N-terminus. The protein was expressed in *E. coli* BL21(DE3) and purified by affinity chromatography on a nickel column (HisTrap HP, GE-Healtcare). Plant eIF4E was prepared in fusion with GST tag attached at the N-terminus in plasmid pGEX4T-1 and expressed in *E. coli* BL21(DE3). The protein was purified on a Glutathione-Sepharose 4B column.

### Interaction of GST-VPg peptides with human eIF4E (ELISA)

The 96-well plates (Nunc) were coated with human eIF4E in 0.1 M carbonate buffer, pH 9.6 (100 ng/well), and incubated overnight at 4°C. Excess eIF4E was removed and the wells were blocked with 5% milk in PBST for 1 hour at 37°C. After milk removal the plates were washed four times (5 min each) with PBST. Two-fold dilutions in cascade of fusion proteins GSTVPg, GSTVPg5, GSTVPg5A1, GSTVPg5A2 and GST alone (2000 – 0.97 nM) were added to the wells and allowed to incubate for 90 min at RT. After removal of proteins excess the wells were washed two times (5 min each) with PBST. Next, the plates were incubated for 1 hour at 37°C with anti-GST-HRP antibody diluted 1:10 000. Excess antibody was removed, the plates were washed three times with PBST and the color reaction was revealed with TMB (Thermo Scientific) and terminated 20 min later with 2 M H_2_SO_4_. The absorbance at 450 nm was measured with the BIO-TEK Synergy HT fluorimeter.

### TAT-VPg peptides

For delivery to human cells in culture, peptides VPg5 and VPg5A2 have been synthesized by Pepscan Presto BV (The Netherlands) as N-terminal TAT fusion peptides, with over 90% of purity.

Tat-VPg5 – NH2-RKKRRQRRR-RKKGKGKGTT VGMGKSSRRF INMYGF-CONH_2_

Tat-VPg5A2 – NH2-RKKRRQRRR-RKKGKGKGTT VGMAKSSRRF INMAGF-CONH_2_

### Effect of Tat-VPg peptides on viability of neoplastic cells in culture

HCT116 cells were maintained in RPMI 40 medium containing 10% fetal bovine serum (FBS), and antibiotics (100 mg/mL streptomycin, 100 U/mL penicillin) at 37°C in a humidified atmosphere of 95% air and 5% CO_2_. They were seeded into 24-well dishes at 1×10^5^ and left overnight. The medium was removed, cells rinsed with PBA and Tat-VPg peptides dissolved in PBS were added for indicated times. The viability was estimated by trypan blue inclusion. The results are expressed as an average of three measurements.

### Level of eIF4E in nuclear and cytoplasmic fraction of HCT116 cells

Before and after treatment with Tat-VPg peptides, HCT116 cells were scraped and fractionated according to REAP method (40). Proteins in cytosol and in the nuclear fraction were fractionated on SDS-PAGE (Mini-protean TGX precast gel from BioRad), transferred to the PVDF membrane using the Trans-Blot Turbo transfer apparatus (BioRad). The eIF4E was revealed with rabbit polyclonal Ab (kind gift of Nahum Sonenberg) at 1:1000 using ECL system (BioRad) and ChemiDoc XRS+ imager (BioRad) for visualisation. Densitometric analysis was done on eIF4E bands using ImageLab software version 4.1 (BioRad).

## Acknowledgement

This work was in part financed by the French Associations Gefluc and Espoir and the Polish National Science Centre. The funders had no role in study design, data collection and interpretation, or the decision to submit the work for publication. The authors have no conflict of interest to declare.

We are indebted to Siergiej Tcherniuk for gift of the plasmid expressing human eIF4E, to N. Sonenberg for gift of the eIF4E antibodies and to L. Strokowskaja and A. Chachulska for gift of the recombinant baculovirus expressing domain II of HCV helicase. RG was the recipient of EMBO and NATO short-term fellowships. This study was in part supported by NATO CLG 982385 grant, ANR 08-EBIO-023 grant, by a Gefluc and Espoir grants and by the Polish Ministry of Sciences and Higher Education grant N302 044 32/3571.

## References

1. Jiang J, Laliberté JF. 2011. The genome-linked protein VPg of plant viruses-a protein with many partners. Curr Opin Virol 1:347–54.

2. Cheng X, Wang A. 2016. The Potyvirus Silencing Suppressor Protein VPg Mediates Degradation of SGS3 via Ubiquitination and Autophagy Pathways. J Virol 91:piie01478–16.

3. Yeam I, Cavatorta JR, Ripoll DR, Kang BC, Jahn MM. (). Functional dissection of naturally occurring amino acid substitutions in eIF4E that confers recessive potyvirus resistance in plants. Plant Cell 2007;19:2913–28.

4. Wang X, Kohalmi SE, Svircev A, Wang A, Sanfaçon H, Tian L. 2013. Silencing of the host factor eIF(iso)4E gene confers plum pox virus resistance in plum. PLoS One 8:e50627.

5. Li H, Kondo H, Kühne T, Shirako Y. 2016. Barley Yellow Mosaic Virus VPg Is the Determinant Protein for Breaking eIF4E-Mediated Recessive Resistance in Barley Plants. Front Plant Sci. 7:1449.

6. Duan H, Richael C, Rommens CM. 2012. Overexpression of the wild potato eIF4E-1 variant Eva1 elicits Potato virus Y resistance in plants silenced for native eIF4E-1. Transgenic Res 21:929–938.

7. Ashby JA, Stevenson CE, Jarvis GE, Lawson DM, Maule AJ. 2011. Structure-based mutational analysis of eIF4E in relation to sbm1 resistance to pea seed-borne mosaic virus in pea. PLoS One 6:e15873.

8. Martínez-Silva AV, Aguirre-Martínez C, Flores-Tinoco CE, Alejandri-Ramírez ND, Dinkova TD. 2012. Translation initiation factor AteIF(iso)4E is involved in selective

9. Joshi B, Cameron A, Jagus R. 2004. Characterization of mammalian eIF4E-family members. Eur J Biochem 271:2189–203.

10. Lejbkowicz F, Goyer C, Darveau A, Neron S, Lemieux R, Sonenberg N. 1992. A fraction of the mRNA 5’ cap-binding protein, eukaryotic initiation factor 4E, localizes to the nucleus. Proc Natl Acad Sci U S A 89:9612–6

11. Strudwick S, Borden KL. 2002. The emerging roles of translation factor eIF4E in the nucleus. Differentiation 70:10–22.

12. Sonenberg N, Gingras AC. 1998. The mRNA 5’ cap-binding protein eIF4E and control of cell growth. Curr Opin Cell Biol 10:268–7513.

13. Sonenberg N. 2008. eIF4E, the mRNA cap-binding protein: from basic discovery to translational research. Biochem Cell Biol 86:178–83

14. Siddiqui N, Sonenberg N. 2015. Signalling to eIF4E in cancer. Biochem Soc Trans 43:763–72

15. Assouline S, Culjkovic B, Cocolakis E, Rousseau C, Beslu N, Amri A, Caplan S, Leber B, Roy DC, Miller WH Jr, Borden KL. 2009. Molecular targeting of the oncogene eIF4E in acute myeloid leukemia (AML): a proof-of-principle clinical trial with ribavirin. Blood 114:257–60.

16. Culjkovic B, Topisirovic I, Skrabanek L, Ruiz-Gutierrez M, Borden KL. 2006. eIF4E is a central node of an RNA regulon that governs cellular proliferation. J Cell Biol 175:415–26.

17. Culjkovic B, Tan K, Orolicki S, Amri A, Meloche S, Borden KL. 2008. The eIF4E RNA regulon promotes the Akt signaling pathway. J Cell Biol 181:51–63.

18. Osborne MJ, Borden KL. 2015. The eukaryotic translation initiation factor eIF4E in the nucleus: taking the road less traveled. Immunol Rev. 263:210–23.

19. Grzela R, Strokovska L, Andrieu JP, Dublet B, Zagorski W, Chroboczek J. 2006. Potyvirus terminal protein VPg, effector of host eukaryotic initiation factor eIF4E. Biochimie 88:887–96.

20. Larsson O, Perlman DM, Fan D, Reilly CS, Peterson M, Dahlgren C, Liang Z, Li S, Polunovsky VA, Wahlestedt C, Bitterman PB. 2006. Apoptosis resistance downstream of eIF4E: posttranscriptional activation of an anti-apoptotic transcript carrying a consensus hairpin structure. Nucleic Acid Res 34:4375–86.

21. Graff JR, Konicek BW, Vincent TM, Lynch RL, Monteith D, Weir SN, Schwier P, Capen A, Goode RL, Dowless MS, Chen Y, Zhang H, Sissons S, Cox K, McNulty AM, Parsons SH, Wang T, Sams L, Geeganage S, Douglass LE, Neubauer BL, Dean NM, Blanchard K, Shou J, Stancato LF, Carter JH, Marcusson EG. 2007. Therapeutic suppression of translation initiation factor eIF4E expression reduces tumor growth without toxicity. J Clin Invest 117:2638–48.

22. Borden KL. 2011. Targeting the oncogene eIF4E in cancer: From the bench to clinical trials. Clin Invest Med 34:E315.

23. Assouline S, Culjkovic-Kraljacic B, Bergeron J, Caplan S, Cocolakis E, Lambert C, Lau CJ, Zahreddine HA, Miller WH Jr, Borden KL. 2015. A phase I trial of ribavirin and low-dose cytarabine for the treatment of relapsed and refractory acute myeloid leukemia with elevated eIF4E. Haematologica 100:e7–9.

24. Rhoads RE. 2009. eIF4E: new family members, new binding partners, new roles. J Biol Chem 284:16711–5.

25. Gingras AC, Raught B, Sonenberg N. 1999. eIF4 initiation factors: effectors of mRNA recruitment to ribosomes and regulators of translation. Ann Rev Biochem 68:913–63.

26. Cohen N, Sharma M, Kentsis A, Perez JM, Strudwick S, Borden KL. 2001. PML RING suppresses oncogenic transformation by reducing the affinity of eIF4E for mRNA. EMBO J 20:4547–59.

27. Volpon L, Osborne MJ, Capul AA, de la Torre JC, Borden KL. 2010. Structural characterization of the Z RING-eIF4E complex reveals a distinct mode of control for eIF4E. Proc Natl Acad Sci U S A 107:5441–6.

28. Maul GG, Negorev D, Bell P, Ishov AM. 2000. Review: properties and assembly mechanisms of ND10, PML bodies, or PODs. J Struct Biol 129:278–87.

29. Topisirovic I, Culjkovic B, Cohen N, Perez JM, Skrabanek L, Borden KL. 2003. The proline-rich homeodomain protein, PRH, is a tissue-specific inhibitor of eIF4E-dependent cyclin D1 mRNA transport and growth. EMBO J 22:689–703.

30. Charron C., Nicolai M., Gallois J.L., Robaglia C., Moury B., Palloix A., Caranta C. 2008. Natural variation and functional analyses provide evidence for co-evolution between plant eIF4E and potyviral VPg. Plant J 54:56–68.

31. Villaescusa JC, Buratti C, Penkov D, Mathiasen L, Planagumà J, Ferretti E, Blasi F. 2009. Cytoplasmic Prep1 interacts with 4EHP inhibiting Hoxb4 translation. PLoS One 4:e5213.

32. Maluccio M, Covey A. 2012. Recent progress in understanding, diagnosing, and treating hepatocellular carcinoma. CA Cancer J Clin 62:394–9.

33. Wang C, Cigliano A, Jiang L, Li X, Fan B, Pilo MG, Liu Y, Gui B, Sini M, Smith JW, Dombrowski F, Calvisi DF, Evert M, Chen X. 2015. 4EBP1/eIF4E and p70S6K/RPS6 axes play critical and distinct roles in hepatocarcinogenesis driven by AKT and N-Ras proto-oncogenes in mice. Hepatology 61:200–13.

34. Zochowska M, Piguet AC, Jemielity J, Kowalska J, Szolajska E, Dufour JF, Chroboczek J. 2015. Virus-like particle-mediated intracellular delivery of mRNA cap analog with in vivo activity against hepatocellular carcinoma. Nanomedicine 11:67–76.

35. Beer S, Zetterberg A, Ihrie RA, McTaggart RA, Yang Q, Bradon N, Arvanitis C, Attardi LD, Feng S, Ruebner B, Cardiff RD, Felsher DW. 2004. Developmental Context Determines Latency of MYC-Induced Tumorigenesis. PLoS Biol 2:e332.

36. Joliot A, Prochiantz A. 2004. Transduction peptides: from technology to physiology. Nature Cell Biology 6:189–196.

37. Jones SW, Christison R, Bundell K, Voyce CJ, Brockbank SM, Newham P, Lindsay MA. 2005. Characterisation of cell-penetrating peptide-mediated peptide delivery. Br J Pharmacol 145:1093–102.

38. Culjkovic B, Borden KL. 2009. Understanding and Targeting the Eukaryotic Translation Initiation Factor eIF4E in Head and Neck Cancer. J Oncol 2009:981679.

39. Fan S, Ramalingam SS, Kauh J, Xu Z, Khuri FR, Sun SY. 2009. Phosphorylated eukaryotic translation initiation factor 4 (eIF4E) is elevated in human cancer tissues. Cancer Biol Ther 8:1463–9.

40. Suzuki K, Bose P, Leong-Quong RY, Fujita DJ, Riabowol K. 2010. REAP: A two minute cell fractionation method. BMC Res Notes 3:294.

41. Schägger H, von Jagow G. 1987. Tricine-sodium dodecyl sulfate-polyacrylamide gel electrophoresis for the separation of proteins in the range from 1 to 100 kDa. Anal Biochem 166:368–79

